# An integrated multi-wavelength SCATTIRSTORM microscope combining TIRFM and IRM modalities for imaging cellulases and other processive enzymes

**DOI:** 10.1101/2021.02.25.432940

**Authors:** Daguan Nong, Zachary K. Haviland, Kate Vasquez Kuntz, Ming Tien, Charles T. Anderson, William O. Hancock

**Author notes:** Corresponding author: William O. Hancock, 122 CBEB, Pennsylvania State University, University Park, PA 16802.

## Abstract

We describe a multimodal SCATTIRSTORM microscope for visualizing processive enzymes moving on immobilized substrates. The instrument combines Interference Reflection Microscopy (IRM) with multi-wavelength Total Internal Reflectance Fluorescence Microscopy (TIRFM). The microscope can localize quantum dots with a precision of 2.8 nm at 100 frames/s, and was used to image the dynamics of the cellulase, Cel7a interacting surface-immobilized cellulose. The instrument, which was built with off-the-shelf components and controlled by custom software, is suitable for tracking other degradative enzymes such as collagenases, as well as motor proteins moving along immobilized tracks.

## 1. Introduction

Cellulases, collagenases, and ribonucleases are all examples of processive enzymes that catalyze degradation of their biopolymer substrates [1-3]. Mechanistic understanding of these enzymes requires visualizing both the individual enzyme molecules, as well as the substrate on which they are acting. Particle tracking by Total Internal Reflectance Fluorescence Microscopy (TIRFM) has been used extensively to study motor proteins moving along their cytoskeletal tracks [4, 5]. These approaches generally require using different fluorophores to separately localize the motors and their tracks. By exciting fluorophores with an evanescent wave generated by total internal reflection, only fluorophores within ∼100 nm of the glass surface are excited, which minimizes background fluorescence and generates high signal-to-noise ratios for single-molecule imaging. By fitting the point spread function of spatially isolated motor proteins, localization down to nanometer spatial precision can be achieved using this approach [6]. This tracking precision is aided by actin filaments or microtubules being easy to polymerize *in vitro* and disperse, which ensures that motors are moving along isolated tracks.

Observing enzymes on more complex substrates such as collagen and cellulose is hindered by the complexity of the underlying substrates. Moreover, these substrates can have thicknesses that extend beyond the evanescent field, complicating interpretation. Furthermore, if both the enzyme and the substrate on which it is acting are fluorescently labeled, crosstalk between fluorescence channels can occur, limiting localization precision. One solution to this problem is to use two different imaging modalities for the mobile enzyme and the surface-immobilized substrate. In the last decade, Interferometric Scattering Microscopy (iSCAT) and Interference Reflection Microscopy (IRM) have been used to image unlabeled actin filaments and microtubules [7-13]. These techniques rely on interference between scattered photons from the sample and reflection at the glass-water interface [7]. They offer high signal-to-noise ratios and, because of the large number of reflected photons, allow for imaging at kHz frame rates [10, 14].

Here, we describe the construction and application of a new SCATTIRSTORM microscope that integrates Interference Reflection Microscopy with multi-wavelength TIRF. The system is built around an open construction commercial microscopy platform, and uses off-the-shelf components. Specific features include both micromirror- and dichroic-based TIRF excitation; integration of the IRM into the dichroic illumination pathway; real-time axial drift correction that stabilizes focus to within 25 nm; and localization precision of 1 nm for a quantum dot at 10 frames/s. Control of lasers, shutters and piezo-stage, as well as image acquisition by the camera is carried out by a custom LabVIEW software package that is freely available to users. As a proof of concept, we used the microscope to image the binding and processive movement of the cellulase Cel7A from *Trichoderma reesei* on bacterial cellulose. The system is flexible and can be adapted for a wide range of biological applications where high-resolution, simultaneous tracking of motile proteins and imaging of complex substrates is needed.

## 2. Methods

### 2.1. Microscope design and assembly

The microscope combines objective-based TIRF through both micromirror and dichroic mirror pathways, together with dichroic-based epifluorescence and IRM (Fig. 1). The microscope is built on an optical table around a Mad City Labs RM21 microscope. Samples are mounted atop a three-axis piezoelectric translation stage (Mad City Labs; USA), Illumination is provided by six lasers: LBX-405-180-CSB-PPA (405 nm), LBX-488-150-CSB-PPA (488nm), LCX-532L-100-CSB-PPA (532 nm), LCX-561L-100-CSB-PPA (561 nm), LBX-647-140-CSB-PPA (647 nm), LBX-785-100-CSB-PPA (785 nm), (Oxxius; France). Each laser beam travels through a f = 25 mm focusing lens (Lf, AL2520-A, Thorlabs; USA), a 25 μm pinhole (Ph, P25Da, Thorlabs; USA), and a f = 250 mm expanding lens (Le, LA1461-A-ML, Thorlabs; USA) to yield a clean and 10-fold expanded beam. The expanded laser beams are combined using five dichroic mirrors (T760lpxr, DM1; T590lpxr, DM2; T545lpxr, DM3; T510lpxr, DM4; T470lpxr, DM5. Chroma; USA) and a broadband mirror, to achieve a co-aligned beam.

**Figure 1:**
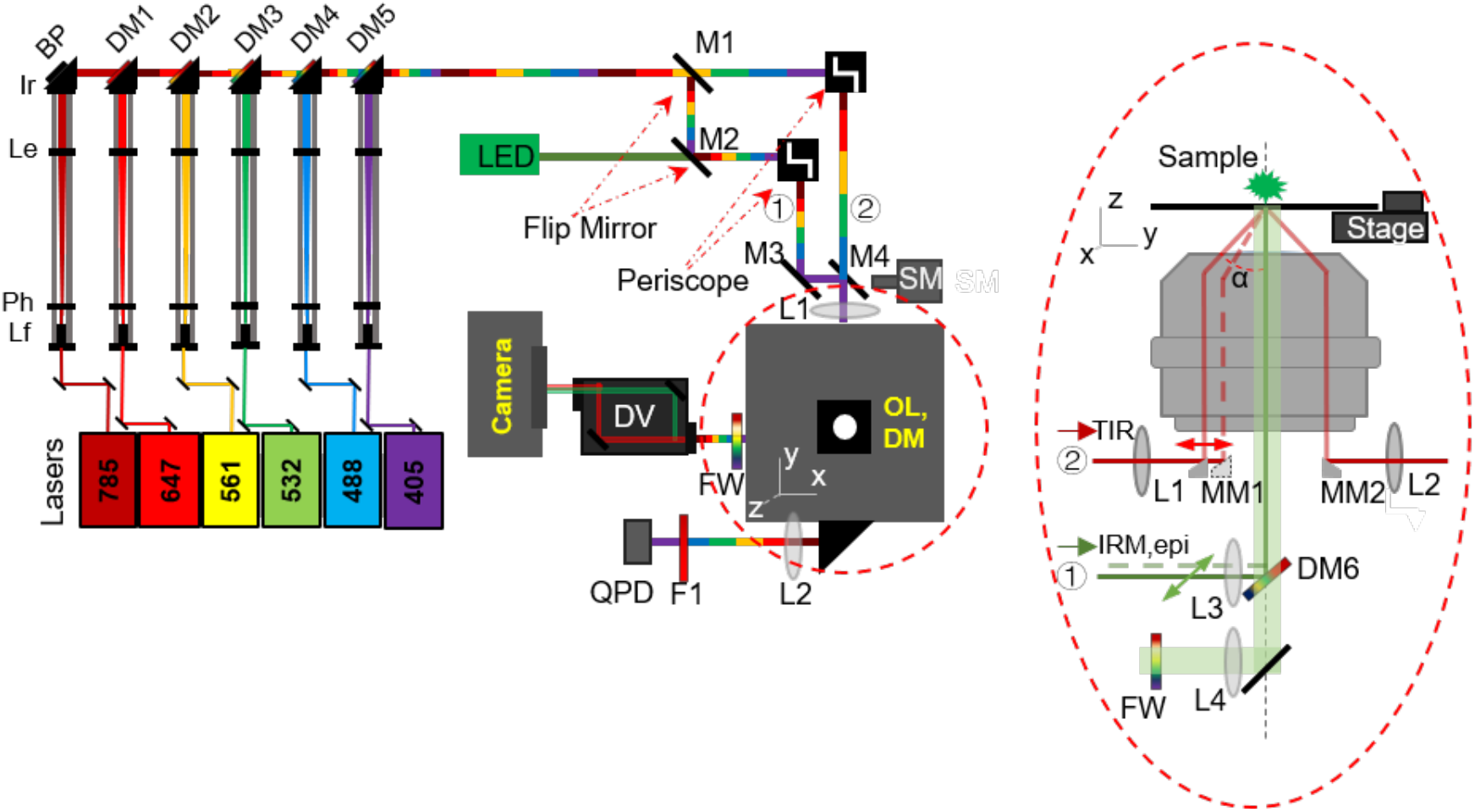
Microscope schematic. Illumination from six lasers at left is combined and collimated by a series of dichroic mirrors. This combined illumination beam is used for micromirror-based illumination (1) or dichroic-based illumination (2), depending on the position of flip mirrors M1 and M2. IRM illumination is supplied by an LED that is reflected off a dichroic mirror below the objective. At right is a detailed view of the objective and associated components, showing micromirrors in the back focal plane and the dichroic mirror (DM6) used for both IRM and dichroic-based TIRF or epi-fluorescence illumination.

The microscope includes two illumination pathways, micromirror and dichroic, that can be switched using a pair of flip mirrors (M1, M2; FM90, Thorlabs; USA). In the micromirror pathway, the beam is raised by a periscope and then passes through a focusing lens (L1, f = 150 mm) mounted on an x-y translation stage (CXY1, Thorlabs; USA). The beam is then reflected off a 2.8-mm ellipsoidal mirror, hereafter termed micromirror (MM1, G54-092, Edmund Optics; USA), positioned in the back focal plane of the objective (APON 60XOTIRF, NA = 1.49, Olympus; Japan). Following total internal reflection at the glass-water interface of the sample, the beam returns through the objective and is reflected by a second micromirror (MM2, G54-092, Edmund Optics; USA) at the back focal plane of the objective, through a focusing lens (L2, f = 250 mm), and onto a quadrant photodiode (QPD, Mad City Labs; USA). The position of this exit beam on the QPD varies with the distance between the objective lens and the sample, and can be used for z-drift correction by the piezo stage.

In the dichroic illumination pathway, the beam is first raised by a periscope, and then reflected by a pair of mirrors (M3 and M4) before passing through a focusing lens (L3, f=150 mm), The beam is then reflected off a dichroic mirror (DM6, ZET488/640m-TRF, Chroma; USA) placed under the objective. M4 is mounted on a translatable stage driven by a stepper motor (SM, Mad City Labs; USA). Translating M4 horizontally results in a corresponding translation of the excitation beam in the back focal plane of the objective, thus enabling switching between epifluorescence and TIRF illumination modes.

The IRM illumination pathway uses a green LED (525 nm, 130 mW; Thorlabs; USA) that is co-aligned with the laser beam before being reflected off the dichroic mirror. Switching between IRM and epi/TIRF modes is accomplished by a flip mirror (M2).

In the imaging pathway, emitted photons collected by the objective first pass through the dichroic mirror (DM6), then through a f = 500 mm focusing lens (L4). Fluorescence emission wavelengths are selected by a filter wheel (CFW6, Thorlabs; USA) and a dual-view system (OptoSplit II, Photometrics; USA) before being focused into the camera (Prime 95B, Photometrics; USA).

### 2.2. Software workflow

All hardware and image acquisition systems are controlled by custom-designed software implemented in LabView in Actor Framework (National Instruments; Austin, TX). The workflow of the main Virtual Instrument (VI) is illustrated in Figure 2. The software was designed to serve as a flexible microscope control platform that can support different hardware modules, and be easily modified to add new extensions based on future imaging needs. The software was designed with four different layers, as follows. 1) The Hardware Layer (HL) handles communication to the specific hardware components. 2) The Hardware Abstraction Layer (HAL) serves as an interface between the hardware and logic layers, and carries out the common functional calls to the hardware (e.g., obtain and set stage position or laser power; set camera exposure and acquire image sequence). 3) The Logic Layer (LL) defines the specific microscope modality and defines specific functions (e.g., switching lasers in a defined sequence). The LL was designed to communicate directly with the HAL and operate without regard to the specific hardware components present. 4) The User Interface Layer (UIL) controls the LL and enables the user to control all aspects of the microscope.

**Figure 2:**
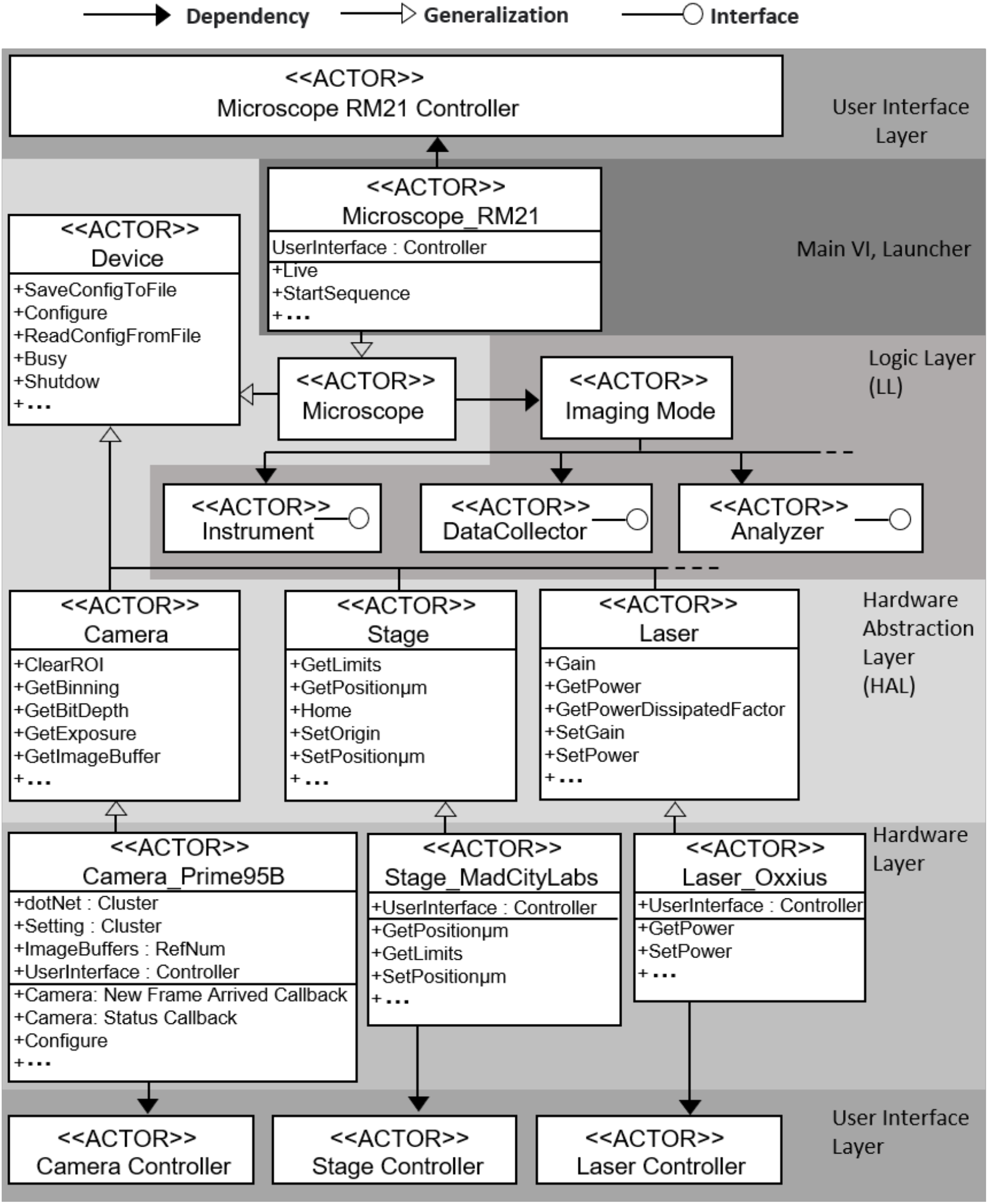
Software workflow for controlling the microscope. See text for details.

Following an object-oriented programing approach, base Actor classes were defined for controlling common hardware elements such as the stage, laser, and camera. Under this framework, hardware components of a similar class are interchangeable. To install new components, such as a new camera, a new Actor derived from the base Actor class is created to directly communicate with the specific hardware component. This strategy minimizes new coding when hardware is replaced or upgraded, or if the software is used in a different microscope. In our microscope, a Camera_Prime95B actor was derived from the Camera base class to control the camera, a Stage_MadCityLab actor was derived from the Stage base class to control the stage and a Laser_Oxxius actor was derived from the Laser base class to control the laser (Figure 2). A TIRF_Lock actor, derived from the Analyser, was created to maintain focus by reading the QPD voltage, comparing it to the lock target voltage, and using a proportional/integrative (PI) controller to move the stage. Images, movies, and hardware status information were handled by the DataCollector actor, and saved for further analysis. The latest version of the software is available on Github (https://github.com/erisir/MMicroscopyAF.git).

### 2.3. Sample preparation and imaging

To test the microscope’s single-molecule imaging capabilities, Acetobacter cellulose and Cel7a enzyme were preprepared as previously described [15]. Cellulose was purified from *G. hansenii* (strain ATCC 23769), sonicated, and microfluidizeed. Cel7a (Sigma) was biotinylated using biotin-NHS (Thermo Scientific, catalog number: 21343). Flow chambers for microscopy were assembled by sandwiching a piece of double-sided tape between a slide and a plasma cleaned coverslip (Fisherbrand; USA). A sample consisting of 20 μL of ∼3 mg/mL cellulose was pipetted into the flow cell and the slide allowed to dry at 80 °C for 5 min until the cellulose fibers were stuck to the coverslip surface. An aliquot of 20 μL of a 1:200 dilution of tetraspeck beads (four excitation/emission peaks of 356/430 nm, 505/515 nm, 560/580 nm, and 660/680 nm; Thermo Scientific) was flowed in and incubated for 5 min. Next, the flow cell was flushed three times with 20 μL of 1 mg/mL bovine serum albumin (EMD Millipore) and incubated for 5 min to block the surface from nonspecific interactions. Cellulose fibers were located using IRM and, after fine-focus adjustments, the TIRF-Lock system was engaged to keep the sample in focus. This was followed by flowing in 20 μL of 2 nM Cel7a labeled with Qdot525 or Qdot655 in 50 mM NaOAc and 5 mM DTT, pH 5.0. Movies of 500 frames of Cel7a reversibly interacting with the immobilized cellulose were captured at 10 frames/s.

## 3. Results and Discussion

### 3.1. TIRF-lock performance

To prevent the sample from drifting out of focus during long data collection periods, a closed-loop focus feedback system was implemented. The distance between the sample and the objective lens was maintained by monitoring the quadrant photodiode (QPD) voltage and moving the piezo-stage to minimize voltage changes (Fig. 3a). The auto-focus system was calibrated by stepping the stage in known increments and recording the QPD output voltage to create a calibration curve (2.26 V/μm; Fig. 3b). Based on this calibration, the z-position of the stage was controlled in the software proportional-integral (PI) control loop. The stability of system was checked by manually applying an external sample displacement of ∼1 μm and observing the system response (Fig. 3c). Upon stage displacement, the QPD voltage in the y-direction dropped immediately, while voltage in the x-direction remained constant, as expected (Fig. 3(c), upper panel). The z-position of the stage was increased in response (Fig. 3(c), bottom panel) until the QPD voltage was returned to its target. After TIRF-lock was established, the standard deviation of the QPD signal in the y-direction was 0.055 V, demonstrating that the system was able to maintain focus with a precision of ± 24 nm.

**Figure 3:**
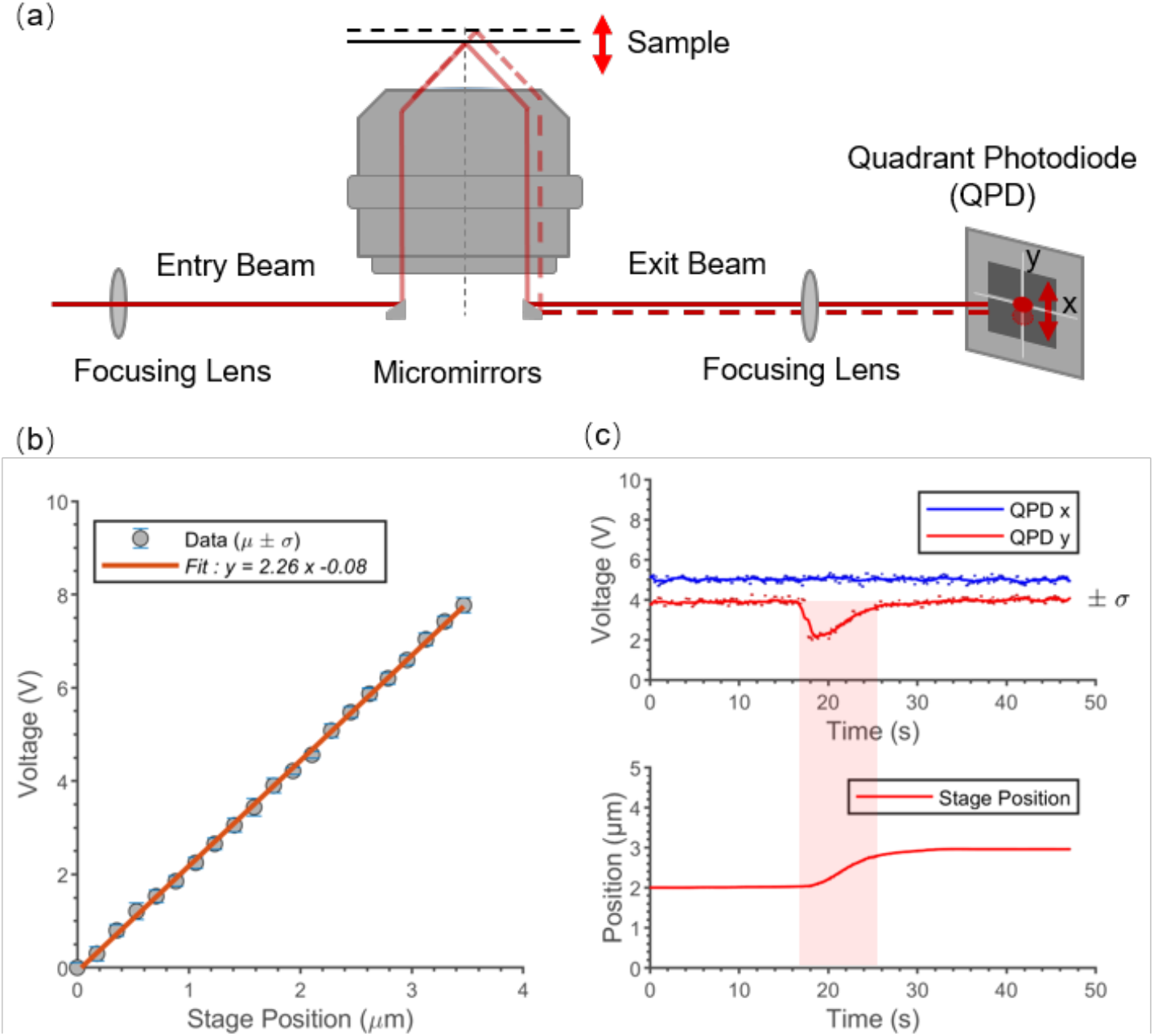
TIRF-lock system. a) Schematic showing how changes in sample height above the objective were detected. An infrared beam (785 nm) collimated with the excitation lasers was totally internally reflected from the glass-water interface of the sample and focused onto the QPD. Changes in sample height caused displacements of the beam that were detected by the QPD. b) Calibration curve of QPD voltage for different z-positions of the stage. c) Step response of the closed-loop feedback system to a ∼1 µm manual step change in the stage height. QPD voltages (top) for vertical (y) and horizontal (x) directions, and stage position (bottom).

### 3.2. Qdot tracking

To track Qdots, we used micromirror TIRF, which minimizes signal loss by dichroic mirrors and facilitates multi-wavelength excitation. Figure 4a depicts a high-constrast quantum dot image (Qdot525, Thermo Scientific, catalog number: Q10143MP), achieved using micromirror excitation by a 405 nm laser.

**Figure 4:**
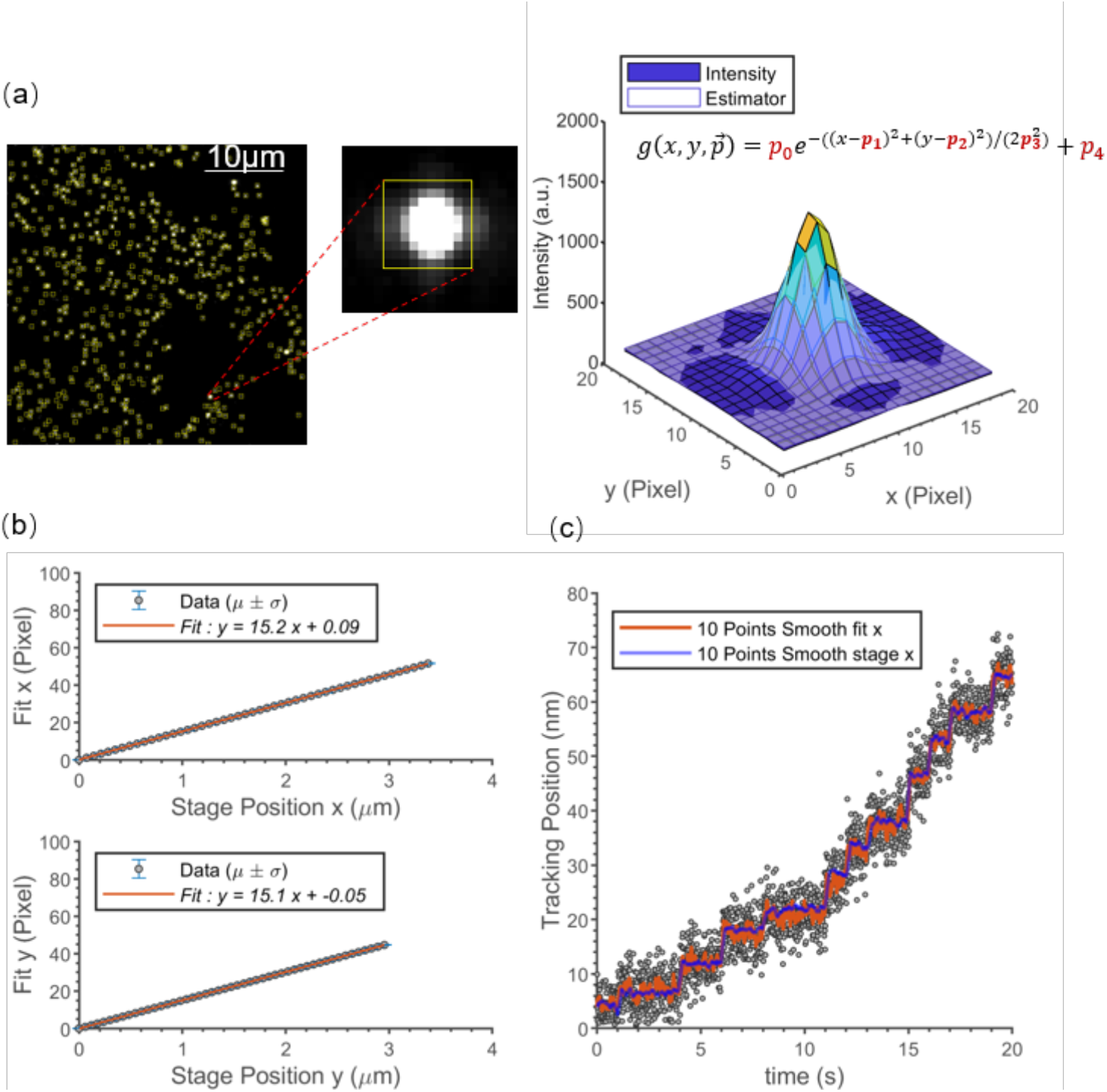
High-resolution tracking of Qdot position. a) Raw images of Qdot-labeled Cel7a bound to immobilized cellulose (left), and image intensity profile of a single Qdot (right). b) Calibration curve to measure pixel size in x (top) and y (bottom). c) Trace of a surface-immobilized Qdot moving due to step changes in stage position. Images were taken at 100 frames/s in a 30 by 30 pixel region of interest. The stage was stepped in random increments between 0 and 20 nm (blue curve), and the Qdot position was calculated by point-spread-function fitting. Gray points are raw position data and the red curve is data smoothed with a 10-point boxcar average. The standard deviation at each step is 2.8 nm for the raw data and 0.87 nm for the 10-point boxcar.

The position of the Qdot was fitted using FIESTA image analysis software [16], which uses Gaussian fitting of the point spread function to achieve sub-pixel position precision (Fig. 4a). To convert pixel size to nm in FIESTA, we measured the representative pixel size by moving the piezoelectric stage in known increments. Figure 4b shows that the line of best fit has a slope of 15.1 pixels/μm (66.2 nm/pixel) in the x and y directions. This value agreed well with our theoretical calculation of 65.98 nm/pixel, calculated by dividing the camera pixel size (11 μm) by the estimated magnification of the microscope (166.7x).

We tested the tracking precision of our system by comparing the step-wise displacements of an immobilized Qdot525 to the step-wise displacements of the piezoelectric stage. The Qdot525 displacements closely matched the stage displacements, and the standard deviation of the plateau regions was 2.8 nm at a frame rate of 100 Hz (Fig. 4(c)).

### 3.3 Simultaneous imaging of Cel7a enzyme and cellulose substrate

Co-visualizing processive enzymes and their substrates is crucial for revealing biological mechanisms. The traditional method of visualizing cellulose is fluorescence microscopy using the cellulose-binding dye Pontamine Fast Scarlet 4B (S4B) or the glucan-binding dye Calcofluor White [17]. However, these dyes can be excited over and emit over broad wavelength ranges, which introduces crosstalk into other fluorescence channels and can degrade the tracking accuracy of enzymes, and the presence of dye molecules on the cellulose surface might interfere with enzyme binding and/or hydrolysis. To achieve label-free imaging of cellulose, we integrated IRM into our multi-wavelength TIRF microscope using a separate imaging pathway. Typically, IRM uses a 50/50 beam splitter in the dichroic position of the standard epifluorescence illumination pathway [18]. Instead, we introduced a separate LED illumination source into the dichroic illumination pathway (Fig. 1). In this configuration, the shared dichroic mirror (Fig. 1; DM6) serves as a ∼ 2:98 beam splitter in the IRM pathway, with a ∼2% reflection and 98% transmission for the 525 nm LED. This arrangement maximizes transmission in the TIRFM channel while maintaining high contrast in the IRM image.

To demonstrate the utility of IRM for imaging cellulose, we compared the intensity profile of an IRM image of label-free cellulose to an intensity profile by TIRF following labeling with the dye S4B. Cellulose was adsorbed to a coverslip and imaged with IRM. The sample was then moved 40 μm out of focus, and a defocused image was taken and used to normalize for any illumination inhomogeneities across the sample field (Fig. 5a, left). Next, 0.5% (w/v) S4B in 50 mM NaOAc was introduced into the sample chamber, incubated for 5 min to allow dye binding, and washed out of the flow cell with 200 μL of 50 mM NaOAc, pH 5.0. Finally, images of the same field were captured by TIRF (Fig. 5a, right). As shown in Figure 5b, the signal intensity profile of IRM was comparable to the TIRF signal, although there was slightly more background signal in IRM due to inhomogeneities in the glass surface and other scattering sources.

**Figure 5.**
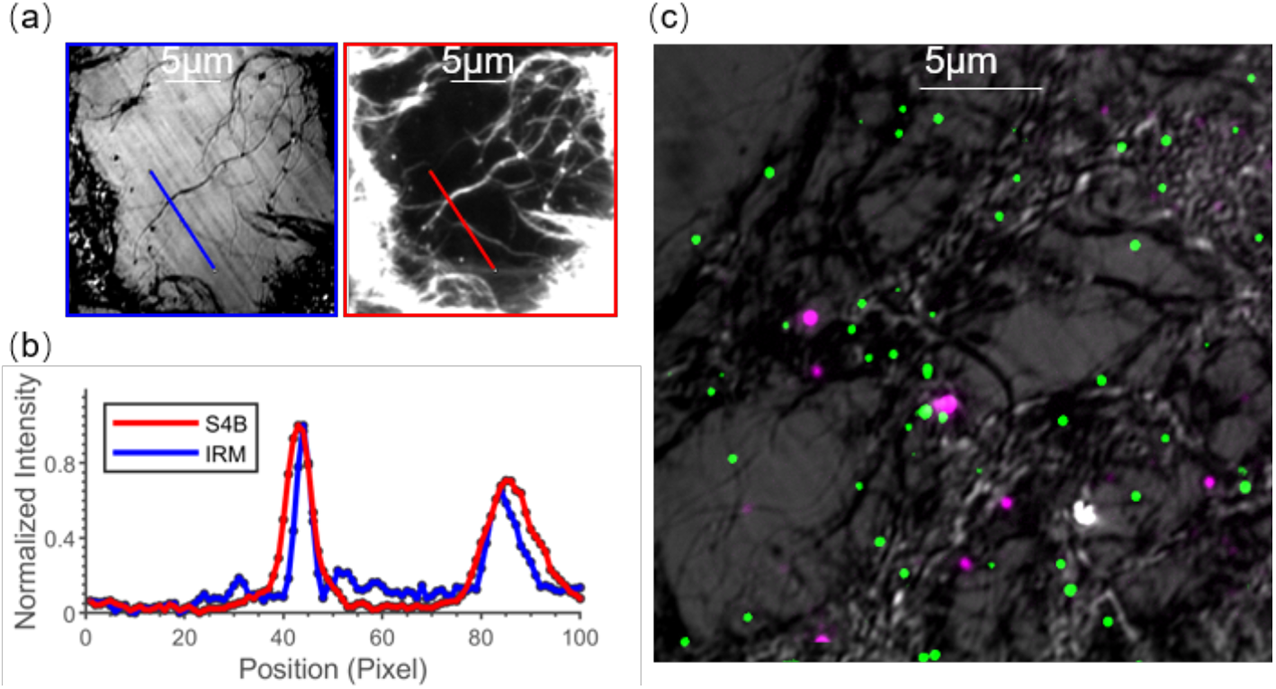
Simultaneous imaging of cellulases and cellulose. a) IRM images of label-free cellulose (left) and TIRFM images of S4B labeled cellulose (right) in the same region. b) Intensity profile comparison of IRM (blue line) and TIRFM (red line). c) Channel overlap of simultaneous imaging of cellulose (black fibers on the gray background), bound Qdot525-labeled-Cel7a (Green), and bound Qdot655-labeled Cel7a (Magenta).

To test the ability of the microscope to simultaneously image cellulose and diverse cellulase enzymes bound to it, we labeled Cel7a with two different quantum dots, Qdot525 and Qdot655. This allowed us to mimic different enzymes binding to the substrate. Figure 5c shows Qdot525-Cel7a (green) and Qdot655-Cel7a (magenta) specifically bound to the unlabeled cellulose fibers (black fibers on the gray background) with high colocalization precision, confirming that the system can be used to image label-free substrate and two different enzyme populations.

### 3.4 Imaging movement of Cel7a cellulase along immobilized cellulose

To demonstrate a practical application of the hybrid IRM-TIRF microscope, we imaged Qdot525-labeled-Cel7a moving along immobilized cellulose (Fig. 6). Image stacks were analyzed by FIESTA to obtain sub-pixel localization of the Qdots. The kinetic behaviors of the molecules were then analyzed and categorized with our custom Matlab software [15]. The movement of one enzyme in x-y coordinates is shown in Figure 6b, and the displacement versus time is shown in Fig. 6(c). The Cel7a molecule moves processively and pauses intermittently. As described in a related study [15], we used this system to analyze 11,116 Cel7a trajectories and categorized their diverse behaviors.

**Figure 6:**
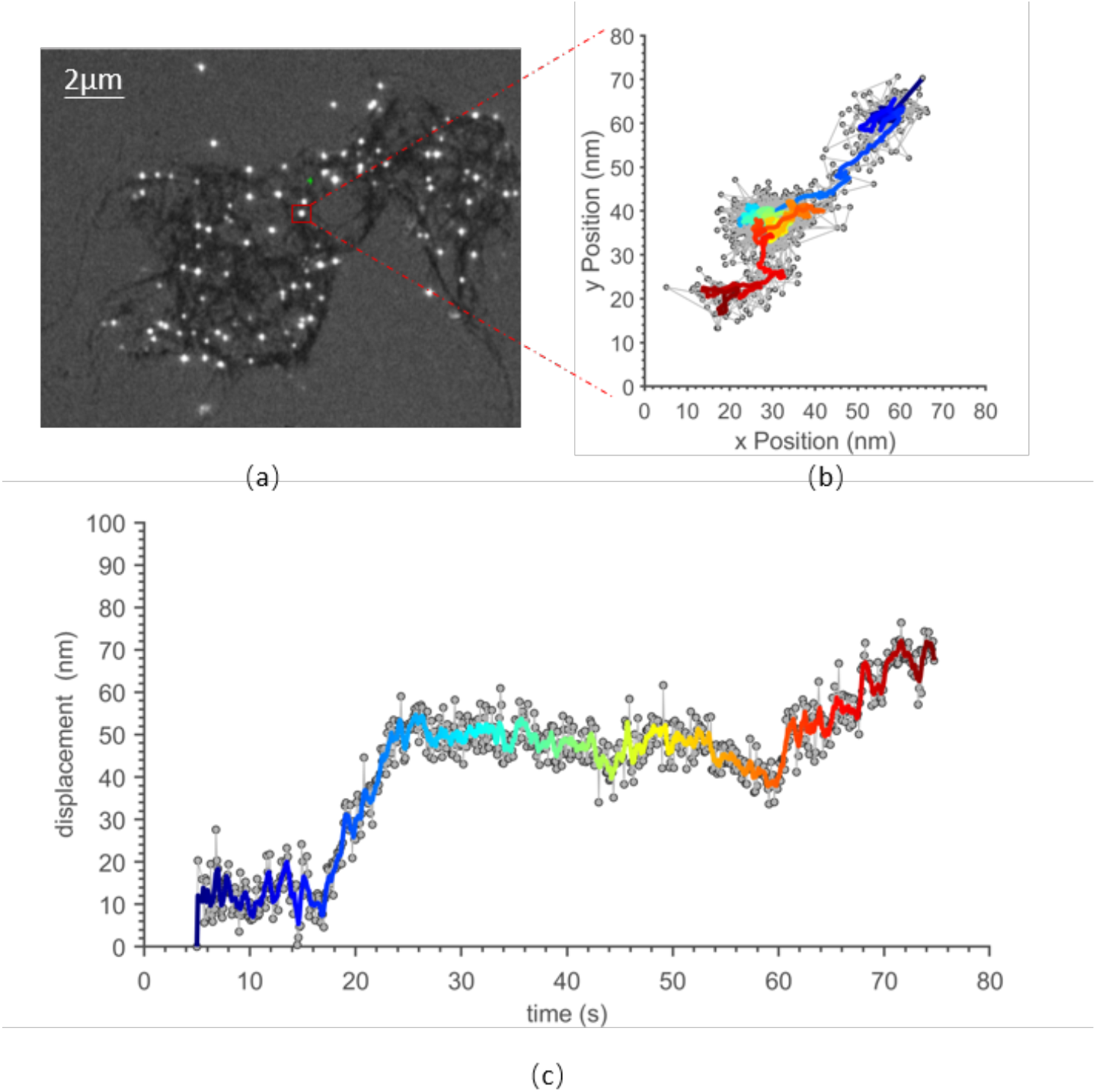
Tracking Cel7a on cellulose. a) Simultaneous imaging of cellulose (black fibers on the gray background) and Qdot525-labeled-Cel7a (bright spots). b) x-y plot of Qdot position, showing Cel7a moving unidirectionally along the cellulose. Gray points are raw data and color curve is data smoothed with a 5-point boxcar. Time is coded by color, from blue to red. c) Displacement from the origin versus time from the same data. Gray points are raw data at 10 fps sampling rate and color curve is data smoothed with a 5-point boxcar, showing the Qdot-labeled Cel7a moving processively along the cellulose with intermittent pauses.

## 4. Conclusion

In the present work we integrated IRM with a multi-wavelength TIRF into a multimodal microscope and developed companion software to control the hardware and image acquisition. Using a closed-loop feedback system, the instrument can stabilize the sample focus within 25 nm. Using micro-mirror TIRF, the system can track Qdots at 100 frames/s and achieve 1 nm positional precision at 10 frames/s. By careful choice of laser lines and filters, two color imaging is relatively simple. By using IRM, substrates can be imaged label-free, which provides significant flexibility. IRM is a relative new technique that has been used to image microtubules with high signal-to-noise [18, 19], and we show here that it is also ideal for imaging immobilized cellulose. We demonstrate that the instrument can image fluorescent Cel7a enzymes degrading cellulose. The instrument has a flexible architecture that also has the capability to expand to iSCAT and STORM microscopy, and these approaches can be applied together to study other processive enzyme-substrate systems such as cytoskeletal motors.

## Acknowledgements

The authors thank Dannielle Gibson and Scott Pflumm for efforts in initiating the cellulase project. The authors also thank the Penn State CSL Behring Fermentation Facility -University Park, PA for assistance with cellulose preparation.

